# The scaffold proteins Tanc1 and Tanc2 are required for myoblast fusion

**DOI:** 10.1101/2022.10.02.510508

**Authors:** Michelle El-Khoury, Andréane Lalonde, David Hipfner, Jean-François Côté

**Author notes:** Corresponding author : Jean-François Côté, IRCM, 110 avenue des Pins Ouest, Montréal (QC) Canada, H2W 1R7.

## Abstract

Myoblast fusion is a crucial step in myogenesis during embryogenesis and adulthood. In *Drosophila*, the scaffold protein Antisocial (Ants)/Rols7 plays is essential in myoblast fusion by connecting the cell adhesion surface proteins to the cytoskeleton. Most molecular pathways governing fusion are evolutionary conserved between mammals and flies, but the relative contributions of Tanc1 and Tanc2, mammalian orthologs of Ants/Rols7, in myoblast fusion have not been established. We used the myoblast C2C12 cell line as a model for differentiation and fusion to assess the contributions of Tanc1 and Tanc2 in fusion. We found that Tanc1 and Tanc2 expressions are not modulated during differentiation, but that both proteins are enriched at the cell cortex in proliferating myoblasts. The knockdown of either Tanc1 or Tanc2 in either of the fusing myoblasts impaired fusion. Notably, the expression of human Tanc1 or Tanc2 restored fusion defects observed in Tanc1- or Tanc2-depleted cells. We found that neither Tanc1 nor Tanc2 could substitute for Ants/Rols7 during *Drosophila* myoblast fusion. We conclude that both Tanc1 and Tanc2 play a role in mammalian myoblast fusion, but there may be some mechanistic differences with the functions of the *Drosophila* orthologous protein.

## Introduction

Myogenesis, or skeletal muscle formation, is a multistep process that involves a cascade of events occurring during embryogenesis and muscle repair^1^. Following the differentiation of the myogenic lineage into myoblast, myoblasts fuse to form multinucleated myotubes. This is a crucial step for the maturation of myofibers. Myoblast fusion has been shown to be essential for proper muscle growth during both embryogenesis and regeneration^2^. Defects in myoblast fusion have been associated with muscular disorders in humans^3-5^.

The study of myoblast fusion during *Drosophila* development has been very powerful to decipher mechanisms underlying cell-cell fusion as many fusion-promoting factors were found to be evolutionarily conserved^1^. Muscle development in *Drosophila* involves the fusion between myoblasts from two genetically distinct populations: Founder cells (FC) and Fusion competent myoblasts (FCM). The initial contact between the two cell types occurs via the interaction of cell surface adhesion proteins specific to each type of myoblast^6^. Dumbfounded /Kin-of-IrreC and Roughest/Irregular-optic-chiasma-C are expressed on the FCs and associate with Sticks and Stones and Hibris found at the surface of FCMs^7-11^. This contact triggers the activation of signaling cascades leading to actin rearrangement^6,11^. Actin nucleation regulators such as Kette, Scar, Wave, Wasp, and Wasp-interacting proteins have been shown to be required in both FCs and FCMs^12-15^.

From the FCM side, the signalling intermediate between the cell adhesion surface proteins and actin nucleation regulators include adaptors (Dock, Drk, Crk), the p21-activated kinases (Paks), the guanine nucleotide exchange factor Myoblast City (Mbc) and its scaffolding protein Elmo and the GTPases of the Rac family^15-23^. In the FC, the principal signaling intermediate is Antisocial (Ants)^24^, also known as Rolling pebbles 7 (Rols7, encoded by the *rols* locus)^25,26^, a scaffold connecting the Dumfounded cell adhesion protein to the actin cytoskeleton. Ultimately, invasive actin-driven podosome-like structures push against the FCs while the FC reply by building a stiffer actin cortex by accumulating MyoII, increasing cortical tension to resist the invasive force until fusion occurs ^13,27,28^.

Although the existence of distinct myogenic lineages in mammals remains unknown, the main cellular events associated with myoblast fusion as well as some key players appear to be conserved. For example, we previously demonstrated that the myoblast fusion function of DOCK1 (ortholog of Mbc) is conserved from flies to mice^29^. Subsequent studies revealed that the conditional inactivation of Rac1 or the nucleating factor N-Wasp in developing mouse myoblasts impairs fusion, further supporting an essential and evolutionarily conserved role for these genes in the fusion process^30,31^. However, *bona fide* fusogenic proteins seem to be specific to vertebrates. Myomaker and Myomerger, also known as Minion and Myomixer were identified as the first muscle specific fusogenic proteins as they can reconstitute fusion when overexpressed together in otherwise non-fusing cells^32-35^. Besides these muscle-specific fusogenic proteins, several broadly expressed cell adhesion proteins and promigratory receptors have been implicated in myoblast fusion^36-39^. The G-protein-coupled receptors Bai1 and Bai3 act upstream of the evolutionarily conserved Elmo/Dock1/Rac1 module for myoblast fusion^40,41^ One essential component for fusion in *Drosophila*, Ants/Rols7, has not been deeply studied in mammalian systems. We therefore focused our interest on its two mammalian orthologs, Tanc1 and Tanc2.

Tanc1 and Tanc2 are scaffold proteins composed of several modular domains known to mediate protein-protein interactions including tetratricopeptide repeat (TPR), ankyrin repeat (ANK), coiled-coil, and C-terminal PSD-95/Dlg/ZO-1 (PDZ) domains, as well as a putative nucleotide triphosphatase (NTPase) domain^42,43^. Tanc1 and Tanc2 expressions patterns and molecular functions have mostly been assessed in the brain^42,44^, but Tanc1 expression has also been detected in various adult tissues including heart, liver, lung and kidney^42^. Their expression differs spatially and temporally suggesting specific functions for each homolog. Tanc1 is highly expressed in the adult brain and Tanc1 knockout mice present impaired spatial memory^44^. Tanc2 is highly expressed during early embryonic stages and Tanc2 knockout mice were embryonically lethal, demonstrating an important role for Tanc2 during embryonic development^44^. Tanc proteins have been shown to interact with postsynaptic density proteins, notably PSD-95^44^ and the glutamate receptor^42^, as well as several signalling molecules associated with the planar cell polarity, Wnt and/or Hippo pathways^43^. According to their predominant roles in the brain, Tanc proteins have been emerging as candidate genes for neurodevelopmental disorders. Notably, Tanc2 mutations have been associated with intellectual disabilities, autism and schizophrenia^45-48^, while Tanc1 mutations have been linked to psychomotor disabilities^49^. Potential roles for Tanc1 in rhabdomyosarcoma, a skeletal muscle cancer, as well as in myoblast fusion have been suggested^50^. However, the contribution of Tanc2 in myoblast fusion has not been characterized.

In this study, we aim to discriminate the relative contribution of Tanc1 and Tanc2 in myoblast fusion using C2C12 cells as a cellular model for differentiation and fusion, and to test whether their functions have been conserved throughout evolution.

## Results

### Tanc1 and Tanc2 are expressed in myoblasts

To study the role of Tanc1 and Tanc2 in mammalian myoblast fusion, we used C2C12, a well-established murine model cell line for myoblast differentiation and fusion. We first verified Tanc1 and Tanc2 expression by Western blot, before and during C2C12 differentiation. Differentiation of C2C12 is induced by placing them in differentiation medium, as shown by the increase of expression of the differentiation marker Myosin heavy chain (Myhc) starting at after 24 h (Fig. 1a). In comparison, Tanc1 and Tanc2 were expressed in proliferating cells and their expression remained constant during the 96 h of differentiation tested (Fig.1a). The relative expression of Tanc1 and Tanc2 was evaluated before and after differentiation by real-time Q-RT-PCR. Tanc1, but not Tanc2, expression increased slightly after 48 h of differentiation (Fig. 1b). We conclude that the protein levels of Tanc1 and Tanc2 remain stable during differentiation despite the small increased in mRNA levels of Tanc1.

**Figure 1.**
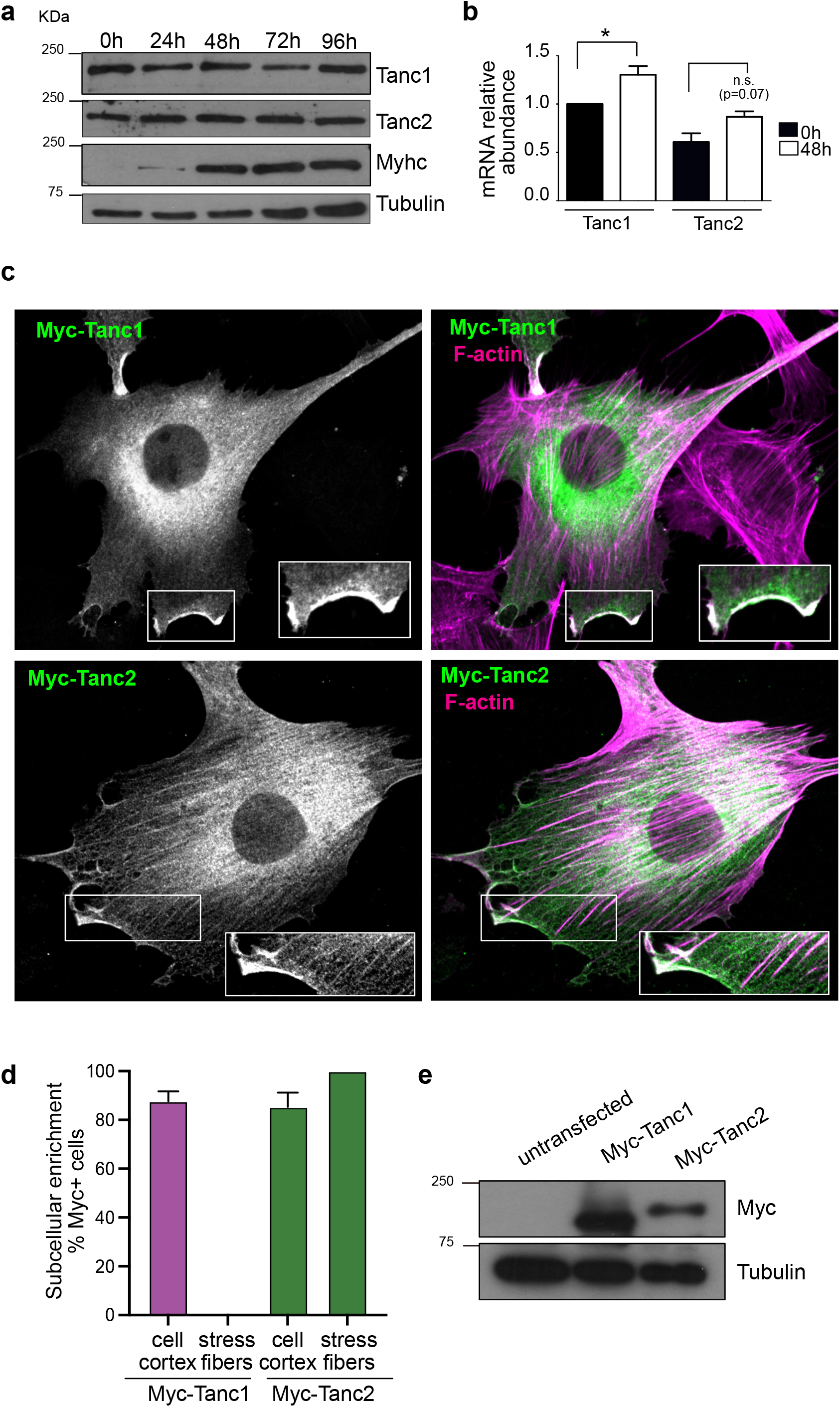
The scaffold proteins Tanc1 and Tanc2 are expressed in myoblasts and are enriched at the cell cortex in proliferating myoblasts. (a-b) Expression of Tanc1 and Tanc2 was measured in differentiating C2C12 cells by Western blot (a) and by real-time Q-RT-PCR (b). Error bars indicate standard deviation. The P value was calculated by one-way ANOVA followed by a Bonferroni test, *P<0.05 (c) Anti-Myc immunostaining of overexpressed Myc-Tanc1 and Myc-Tanc2 in proliferating C2C12 cells. Enlargement (insets) shows the enrichment of Tanc1 and Tanc2 at the cell cortex. (d) Quantification of the cellular localizations of Tanc1 and Tanc2 at the cell cortex or stress fibers in proliferating cells. Data are the mean +/-SD of three independent experiments. (e) Western blot validation of the overexpression of Myc-Tanc1 and Myc-Tanc2. Every experiment was replicated three times. Original blots are presented in supplementary Fig S5.

Based on the hypothesis that Tanc1/2 partake in fusion, we expected them to be recruited at the cell cortex, like their *Drosophila* ortholog Ants/Rols7^24,25^. Localization of Tanc1 and Tanc2 was assessed by immunofluorescence by overexpressing Myc-tagged Tanc1 and Tanc2 proteins (Fig. 1c-e). In exponentially growing cells, both Tanc1 and Tanc2 were localized in the cytoplasm with a fraction enriched at the cellular cortex in more than 75% of the expressing cells (Fig.1d). In addition, Tanc2, but not Tanc1, was detectable along actin stress fibers (Fig. 1c-d). Collectively these data show that Tanc1 and Tanc2 are both expressed in myoblast cells during proliferation and differentiation of C2C12 and are both specifically enriched at the cortex of the myoblasts during proliferation.

### Tanc1 and Tanc2 are not involved in the differentiation of myoblasts

The enrichment of both Tanc1 and Tanc2 at the cell cortex in myoblasts position them as potential scaffold proteins involved in membrane fusion. To verify the potential contributions of Tanc1 and Tanc2 in myoblast differentiation and fusion, we generated C2C12 stable cell lines that express small hairpin (sh)RNA targeting either murine Tanc1 or Tanc2 using a retroviral expression system where GFP is used as a selection marker. The specificity and efficiency of the knockdown of Tanc1 and Tanc2 were validated by real-time Q-RT-PCR (Fig. 2a-b) and Western blot (Fig. 2c). Since myoblasts need to differentiate before fusion, alteration in the differentiation program could manifest as a fusion defect. Consequently, we wanted to rule out the contribution of Tanc1 and Tanc2 in myoblast differentiation, before testing the shRNA system in myoblast fusion. To assess differentiation, the expression of classical myogenic regulatory factors was monitored by real-time Q-RT-PCR and Western blot. As expected, the induction of the expression of Myod1 (Fig. 2d), Myog (Fig. 2e) and Myh4 (Fig. 2f) at the mRNA level was not affected by the depletion of Tanc1 and Tanc2, excluding a central role for these genes in differentiation. Also, the expression of the mRNAs of Mymk (Fig. 2g) and Mymx (Fig. 2h), encoding fusiogenic proteins, were normally induced in Tanc1/2 depleted cells. The expression levels of Myosin heavy chain (Myhc), Myogenin and Myoblast determination protein 1 (Myod) proteins were not altered by depletion of either Tanc1 or Tanc2 (Fig. 2i). These results exclude a central role of Tanc1 and Tanc2 in myoblast differentiation.

**Figure 2.**
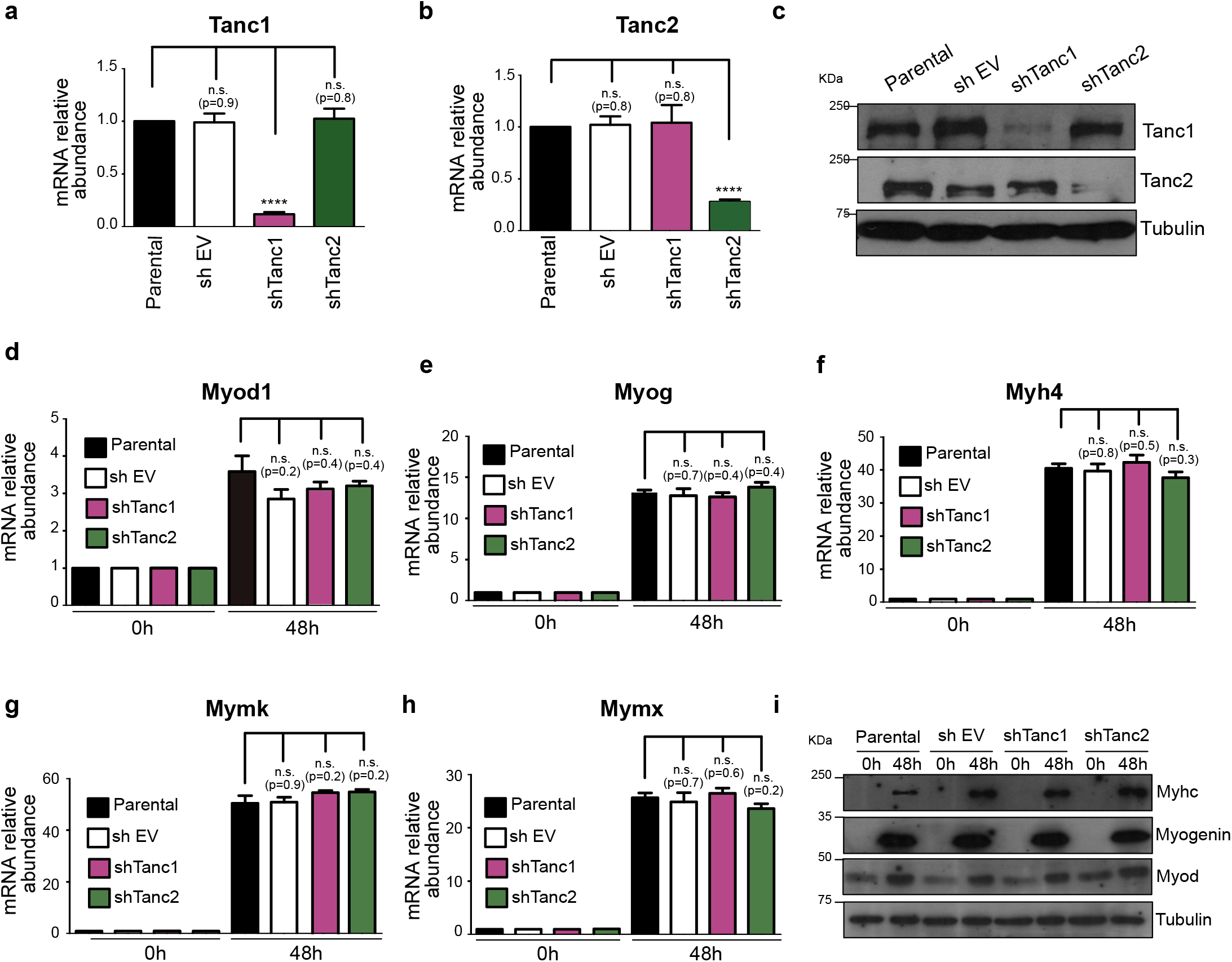
Tanc1 and Tanc2 are not involved in the differentiation of myoblasts. (a-c) The specific knockdown of Tanc1 and Tanc2 in C2C12 was validated by real-time Q-RT-PCR and Western blot. (d-h) Real-time Q-RT-PCR was performed with differentiation and fusion markers to study the impact of *Tanc1* and *Tanc2* knockdowns on the expression of *Myod1* (d), *Myog* (e) *Myhc4* (f), *Mymk* (g) and *Mymx* (h). Error bars indicate standard deviation. The P value was calculated by one-way ANOVA followed by a Bonferroni test. (i) The expression of the indicated differentiation markers was validated by Western Blot. Every experiment was replicated three times. Original blots are presented in supplementary Fig S6.

### Tanc1 and Tanc2 are both important for myoblast fusion

After validating the establishment of the differentiation program in Tanc1 or Tanc2 knockdown myoblasts, we next wanted to specifically assess the role of Tanc1 and Tanc2 in myoblast fusion. ShTanc1 and shTanc2 cells were subjected to differentiation assays and the percentage of multinucleated Myhc+ myoblasts were evaluated as a read-out of myoblast fusion (Fig. 3a-b). In the parental and empty vector-infected cells, most of the Myhc+ fibers are multinucleated (Fig. 3a-b) after 48 h of differentiation. We calculated the fusion index, i.e. the % of nuclei found in fibers with 3 or more nuclei over the total number of nuclei. In comparison, the knockdown of either Tanc1 or Tanc2 completely blocked myoblast fusion as shown by the absence of multinucleated Myhc+ fibers of 3 nuclei or more in shTanc1 and shTanc2 cells. The same results were obtained using different shRNA target sequences against Tanc1 and Tanc2 (Supplementary Fig. S1). Notably, impaired fusion was not due to a delay since Tanc1 and Tanc2 knockdown cells were unable to fuse even after 96 h of differentiation (Supplementary Fig. S2). The knockdown of Tanc1 and Tanc2 also impaired myoblast fusion in Sol8 myoblast cell line (Supplementary Fig. S3). These results show that Tanc1 and Tanc2 are essential players in the fusion of myoblast cell lines.

**Figure 3.**
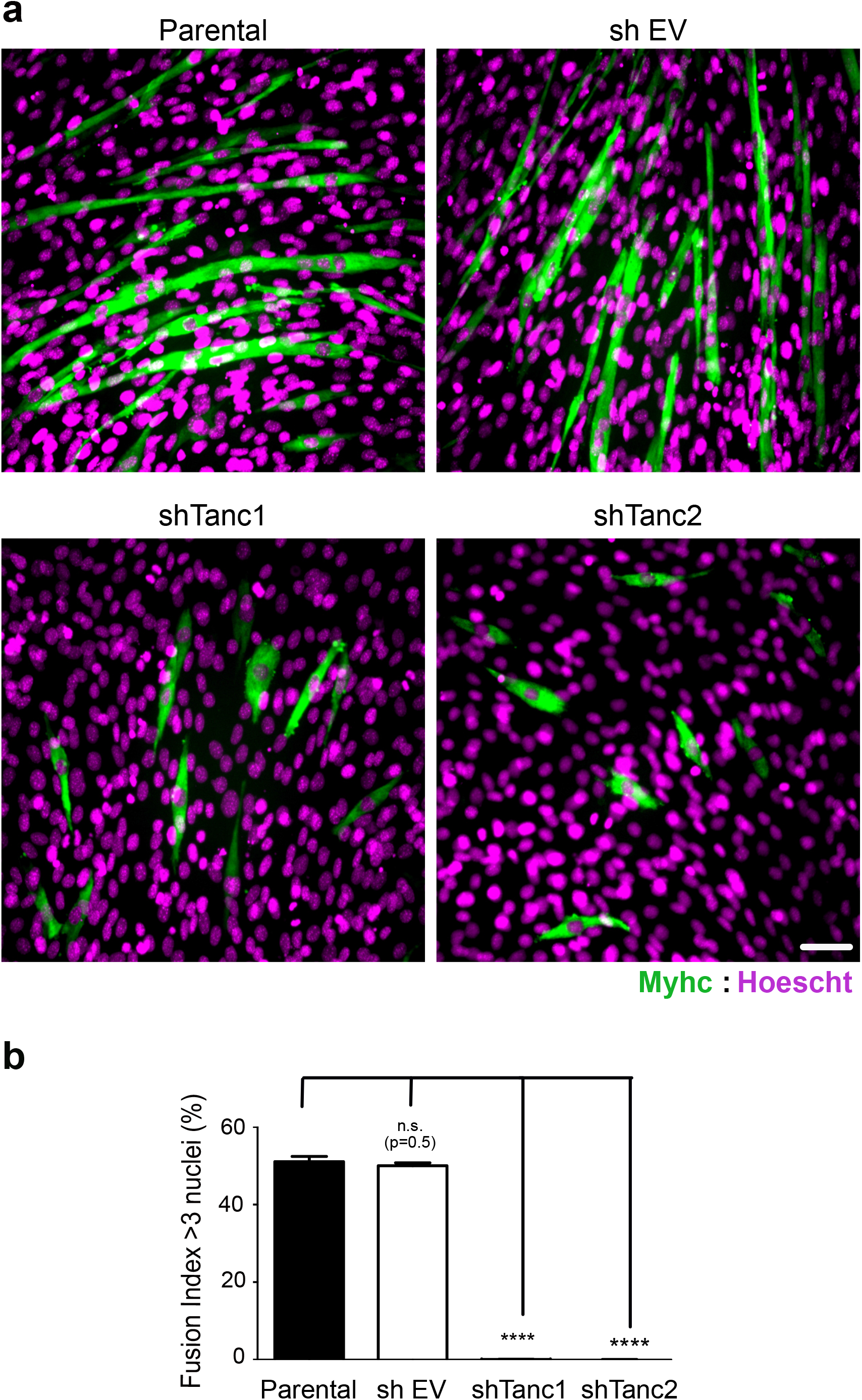
Tanc1 and Tanc2 are both required for myoblast fusion. (a) Parental, shTanc1 and shTanc2 myoblast lines were stained with Myhc and Hoechst after 48 h of differentiation to specifically assess the contributions of Tanc1 and Tanc2 to fusion. No multinucleated fibers (3 nuclei or more) were detected in shTanc1 and shTanc2 cells. Scale bars, 50 μm (b) Fusion index (number of nuclei incorporated in Myhc+ fibers of 3 nuclei or more over the total number of nuclei) was calculated on the conditions presented in (a). Data represent mean +/- SD of 3 independent experiments. The P value was calculated by one-way ANOVA followed by a Bonferroni test, ****P<0.0001.

To confirm that the absence of multinucleated fibers was caused by the absence of Tanc1 or Tanc2 proteins, rescue experiments were performed by overexpressing Myc-tagged human Tanc1 or Tanc2 protein (Fig. 4 a-d). The overexpression of Myc-tagged human Tanc1 in shTanc1 and of Myc-tagged human Tanc2 in shTanc2 rescued the fusion defects seen in shTanc1 and shTanc2 cells, respectively (Fig. 4a-c). Interestingly, the expression of human Myc-Tanc1 in Tanc2 shRNA cells and of human Myc–Tanc2 in Tanc1 shRNA cells fully restored the formation of multinucleated Myhc+ cells (Fig. 4a-c). The expression levels of the endogenous and exogenous Tanc1 and Tanc2 proteins was verified by western blot and quantified (Fig. 4c-d). Collectively, these data suggest that Tanc1/2 might carry out similar essential functions and that a threshold level of their expression is required for efficient myoblast fusion.

**Figure 4.**
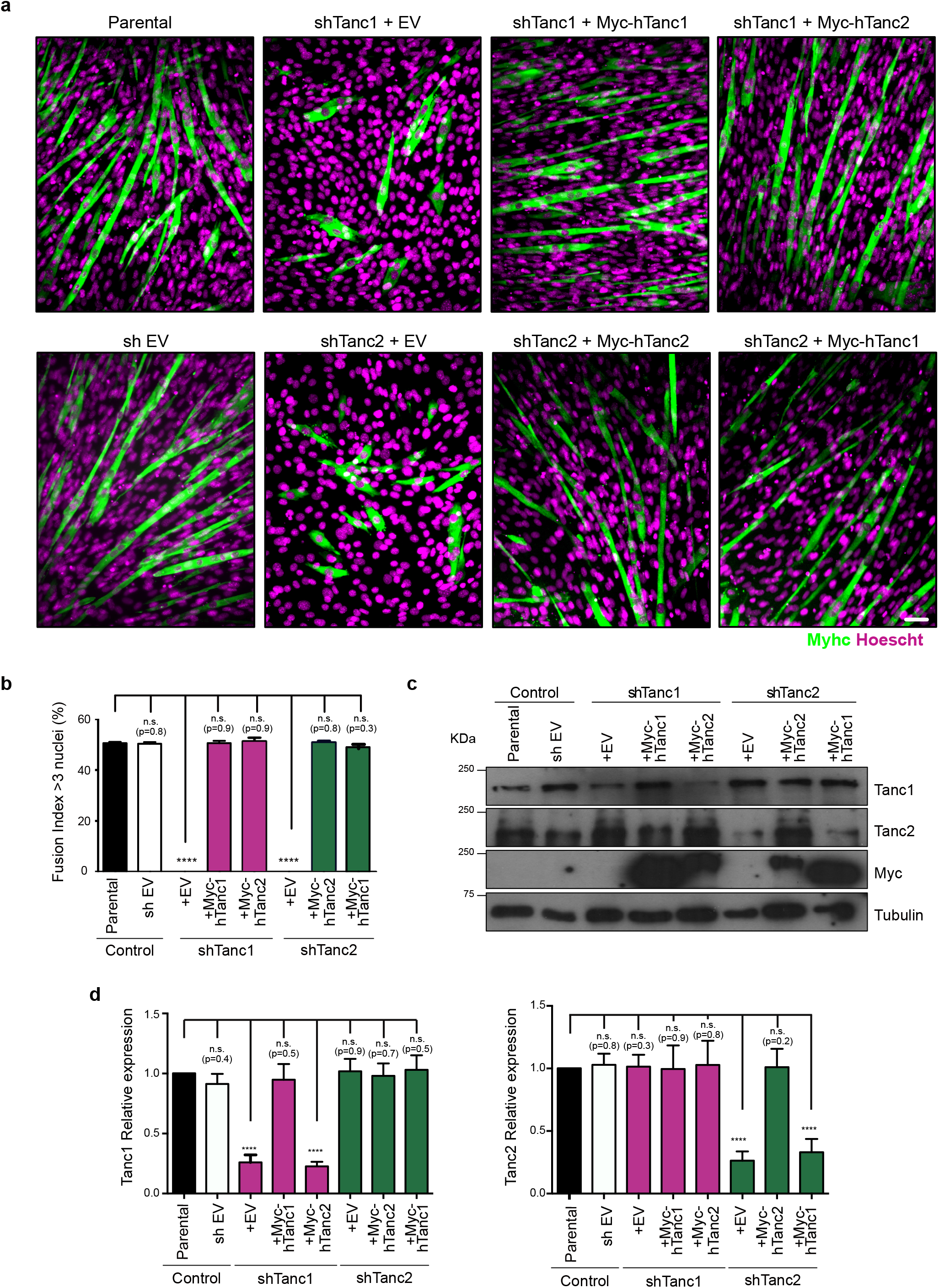
Rescue experiment reveals that Tanc1 and Tanc2 play overlapping roles in myoblast fusion. (a) ShTanc1 and shTanc2 phenotypes were rescued by overexpressing Myc-Tanc1 and Myc-Tanc2 in shTanc1 and shTanc2 cells, respectively. Myc-Tanc1 and Myc-Tanc2 were also expressed in shTanc2 and shTanc1 cells, respectively. The impaired fusion phenotypes were successfully rescued in all conditions. Fusion index (number of nuclei incorporated in Myhc+ fibers of 3 nuclei or more over the total number of nuclei) was calculated on the conditions presented in (a). (c) Validation of the of Myc-Tanc1 and Myc-Tanc2 proteins in the indicated myoblast cells by Western blot. Original blots are presented in supplementary Fig S7. (d) Quantification of the protein levels of endogenous and exogenous Tanc1 and Tanc2 in the indicated myoblast cell lines. Error bars indicate standard deviation. The P value was calculated by one-way ANOVA followed by a Bonferroni test, ****P<0.0001. Scale bars, 50 μm. Every experiment was replicated three times.

### Tanc1 and Tanc2 are required on both cells for efficient myoblast fusion

Since Rols7 was shown to be needed only in one of the fusing cells^24,26^, we investigated whether Tanc1 and Tanc2 are needed in one or both fusing myoblasts. shTanc1 and shTanc2 cells were labeled with a green lipid dye and mixed in a 1:1 ratio with parental cells labeled with a red lipid dye. After 24 h, cells were subjected to a differentiation assay for 48 h. A mix of green and red labeled parental cells as well as a mix of green labeled shTanc1 and red labeled shTanc2 were used as controls. In the mix between green and red labeled parental cells, yellow myotubes were mainly observed, indicating fusion between red and green populations (Fig. 5a). However, when green labeled shTanc1 cells were mixed with red labeled parental cells, red multi-nucleated fibers and green mono-nucleated cells were mainly detected. This indicates that no fusion occurred between the parental and the shTanc1 cells (Fig. 5b). The same phenotype was observed in the mix between green labeled shTanc2 and red labeled parental cells (Fig. 5c). In the mix between green labeled shTanc1 and red labeled shTanc2, red and green mono-nucleated cells were mainly detected (Fig. 5d). Collectively, these results suggest that Tanc1 and Tanc2 are each needed on both myoblasts for proper fusion in a vertebrate myoblast model.

**Figure 5.**
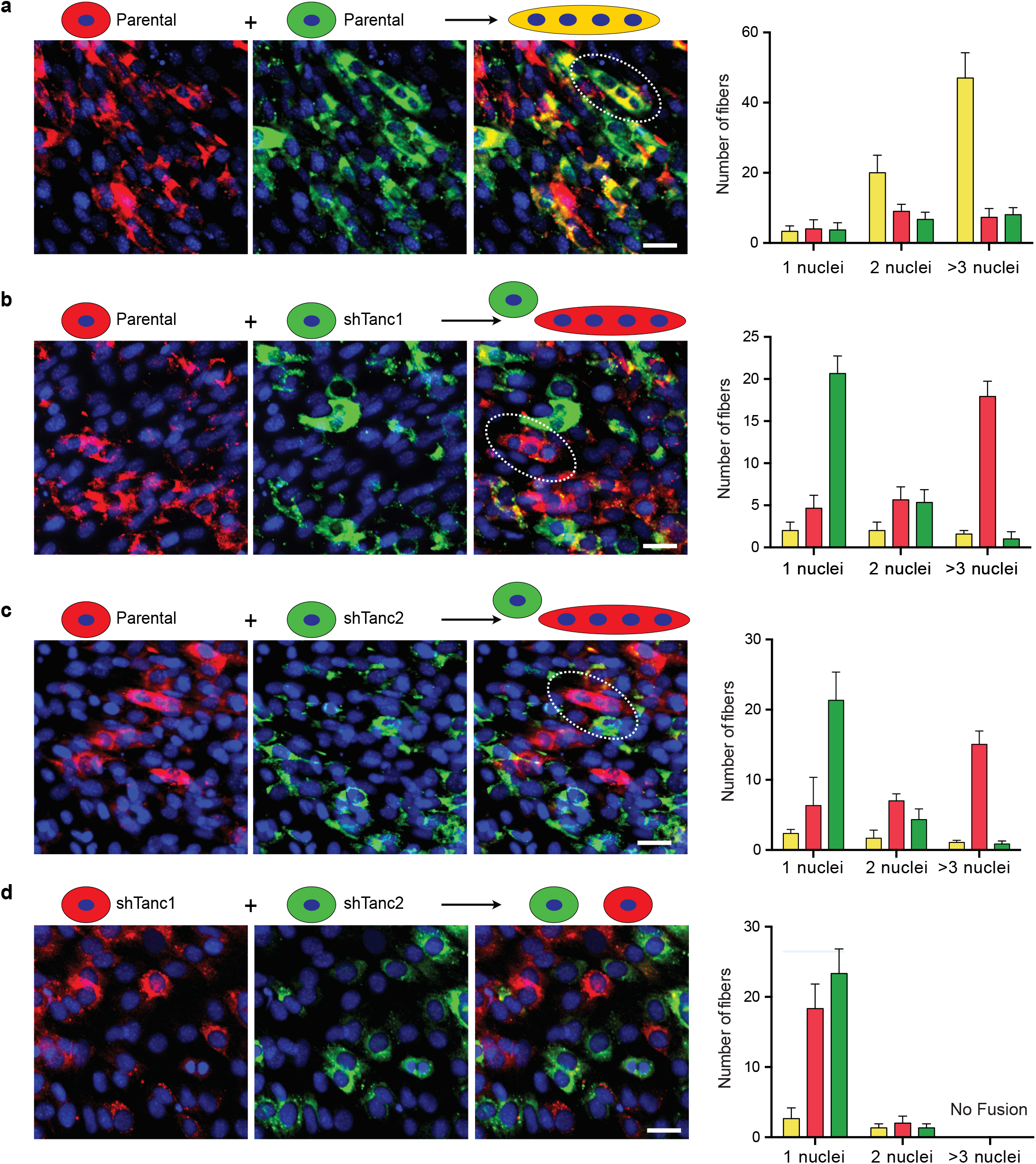
Tanc1 and Tanc2 are needed on both fusing cells. (a) Parental cells were stained with either green or red lipid dyes and mixed in a 1:1 ratio. 24 h following plating, cells were induced to differentiate for 48h. Yellow multi-nucleated fibers are observed because of fusion between green and red cells. (b) Mixing of red parental cells with green shTanc1 mainly resulted in red multi-nucleated fibers and green mono-nucleated cells. (c) Mixing red parental cells with green shTanc2 cells resulted in red multi-nucleated fibers and green mono-nucleated cells. (d) Mixing of green shTanc1 cells with red shTanc2 cells generated only green and red mono-nucleated cells indicating impaired fusion. Scale bars, 50 μm. Data are the mean +/- SD of three independent experiments.

### Do Tanc1/2 and Rols7 have conserved functions in myoblast fusion?

Our results suggested that Tanc1 and Tanc2 may play a similar function in mammalian myoblast fusion as Ants/Rols7 does in *Drosophila*, despite the absence of a conserved fusion receptor acting upstream. Both mammalian Tanc proteins show substantial sequence conservation with Rols7. The three proteins share 38% identity and 70% similarity across the conserved core of the protein spanning from the putative NTPase domain through the ANK and TPR regions (Fig. 6a) ^43^. We therefore decided to test if Tanc1 or Tanc2 could substitute functionally for Ants/Rols7 in Drosophila myoblast fusion.

**Figure 6.**
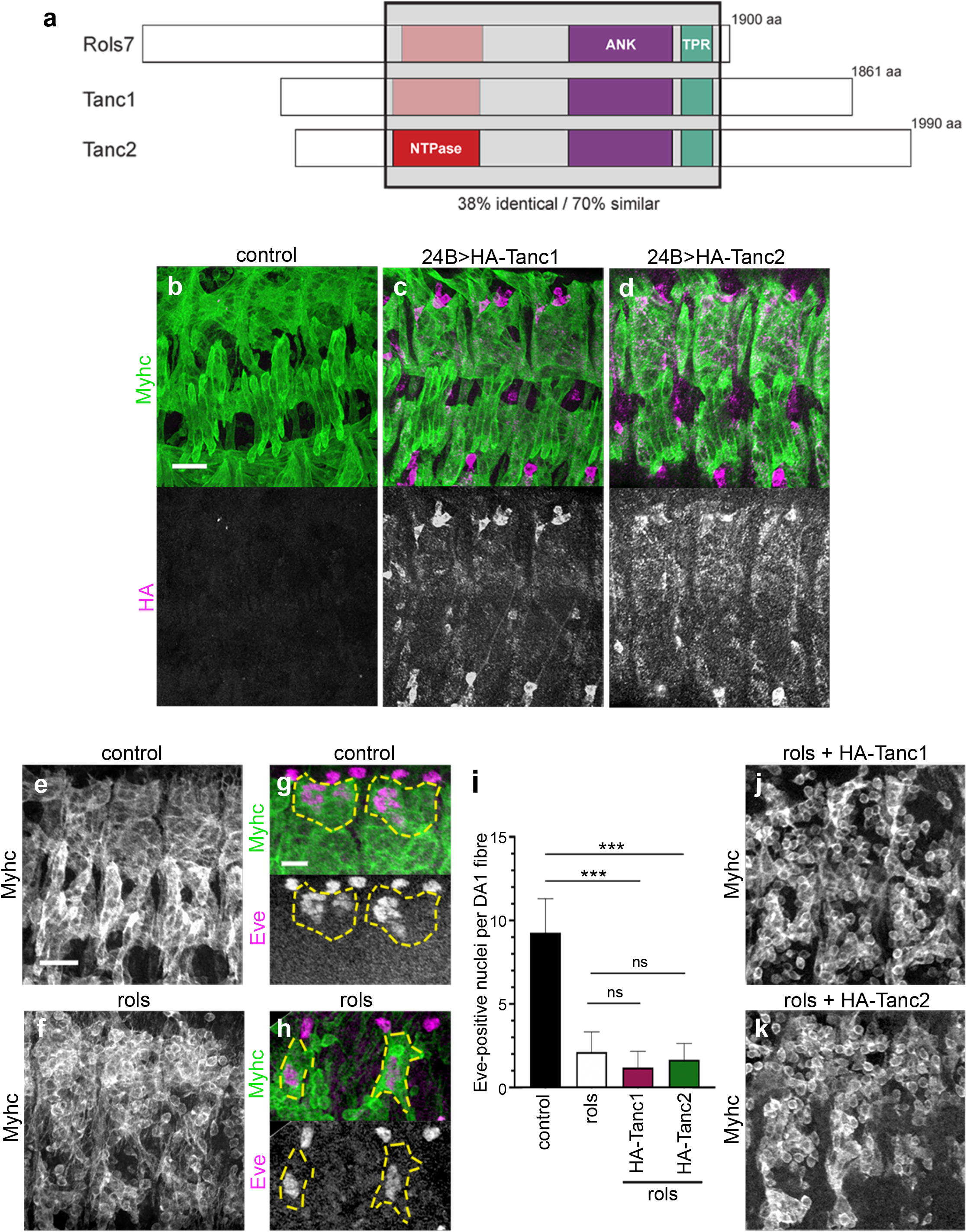
Neither Tanc1 nor Tanc2 can substitute for Drosophila Rol7. (a) Tanc1 and Tanc2 show substantial sequence homology (38% identity, 70% similarity) to Drosophila Ants/Rols7 through the conserved core that includes a putative P-loop nucleoside triphosphate hydrolase (NTPase) domain (predicted in Tanc2) through to the Ankyrin (ANK) and tetracopeptide repeat (TPR) domains. (b-d) Detection of *24B-Gal4*-driven HA-Tanc1 and HA-Tanc2 expression in the developing musculature of stage 15 embryos, marked by Myhc staining. Scale bar: 20 μm. (e-h) Compared to wild-type control (e,g), the myoblast fusion defect in stage 14 *rols* mutant embryos (f,h) is evident from the decreased size of muscle fibres, presence of many rounded unfused myoblasts, and reduced number of Eve-positive muclei in the DA1 fibre. Scale bars: 20 μm (e-f); 10 μm (g,h). (i) Quantification of the number of Eve-positive DA1 nuclei in the indicated gentoypes. *t*-test of statistical significance: ***, *p*<.001; *ns*, not significant. (j-k) Stage 14 *rols* mutant embryos with *24B-Gal4*-driven expression of HA-Tanc1 or HA-Tanc2 in myoblasts show a similar fusion defect as *rols* mutants. Scale bar: 20 μm. Genotypes: *w;24B-Gal4/+* (b); *w;UAS-HA-Tanc1/+;24B-Gal4/+* (c); *w;UAS-HA-Tanc2/+;24B-Gal4/+* (d); *w*^*1118*^ (e,g); *w;rols*^*T192*^*/rols*^*T192*^ (f,h);*w;UAS-HA-Tanc1/+;24B-Gal4,rols*^*T627*^*/rols*^*T192*^(j);*w;UAS-HA-Tanc2/+;24BGal4,rols*^*T627*^*/rols*^*T192*^ (k). The rescue experiment was performed twice, n=12 embryos total per condition.

We generated transgenic strains for expressing N-terminally HA-tagged Tanc1 and Tanc2 *in vivo* under the control of UAS sequences. Both proteins were detectable in the developing musculature (marked by Myhc staining) when expressed in wild-type embryos using the muscle-specific *24B-Gal4* driver (Fig. 6b-d). Expression was already detectable in individual myoblasts at stage 13, when fusion is ongoing (Supplementary Fig. S4). Neither overexpression condition interfered with normal development (data not shown). *rols* mutant embryos display a characteristic muscle fusion defect characterised by fewer irregularly shaped miniature muscles containing a reduced number of nuclei, surrounded by many rounded unfused myoblasts (Fig, 6e-f)^24,25^. Quantification of this defect by counting the number of Even-skipped (Eve)-positive nuclei in the DA1 muscle fibre at stage 15 yielded an average of 9.4 nuclei in wild-type and 2.5 nuclei in *rols* mutants (Fig. 6g-i), consistent with previously reported values^25,26^. *24B-Gal4*-driven expression of N-terminally HA-tagged Ants/Rols7 was previously shown to fully rescue the myoblast fusion defects in *rols* mutants^25^. However, Tanc1 and Tanc2-expressing *rols* mutant embryos still showed prominent unfused myoblasts (Fig. 6j-k), and there was no increase in the number of Eve-positive nuclei in DA1 fibers (Fig. 6i). This suggests that Tanc1 and Tanc2 do not have the same precise function in fusion as Ants/Rols7, or that sequence divergence prevents them from functioning properly in *Drosophila*.

## Discussion

Several fusion promoting factors are evolutionarily conserved between *Drosophila* and vertebrates. For example, the Elmo/Dock1/Rac1 pathway that relays signals from the adhesion proteins to the cytoskeleton in the fusion-competent myoblasts in *Drosophila* is required for myoblast fusion in vertebrates by transmitting signals from Bai1 and Bai3 cell surface proteins^40,41,51^. So far, the conservation of the signalling pathways that relays the signal from the adhesion surface proteins of the *Drosophila* founder cells to the cytoskeleton, orchestrated by the scaffold protein Ants/Rols7, remained unclear. Ants/Rols mammalian ortholog Tanc1 was shown to be involved in myoblast fusion^50^, but the role of the second ortholog, Tanc2, has never been assessed. Here we show that the two mammalian orthologs of Ants/Rols7, Tanc1 and Tanc2, are essential players during myoblast fusion in mammalian myoblasts. Using the murine C2C12 myoblasts as a model for fusion, we report that Tanc1 and Tanc2 are expressed in myoblasts and localized at the cell cortex. The knockdown of either Tanc1 or Tanc2 impaired the formation of multinucleated fibers by specifically blocking myoblast fusion implicating a role for these proteins in cell-cell fusion.

To claim Tanc1 and Tanc2 as evolutionarily conserved functional orthologs of Ants/Rols7, a crosstalk with the actin cytoskeleton still needs to be demonstrated. Supporting this hypothesis, Tanc1/2 have been shown to work as postsynaptic scaffolds by forming a multiprotein complex with several postsynaptic density (PSD) proteins in neurons^42^. The post synaptic membrane is composed of cell adhesion molecules bound to various scaffold proteins that connect to the cytoskeleton. In this study, we report that overexpressed Tanc1 and Tanc2 are enriched at the cell cortex together with polymerized actin as well as along actin stress fibers in the case of Tanc2. This suggests a potential interplay between Tanc1/2 and polymerized actin and support the hypothesis that Tanc1/2 could play a scaffolding role that organizes proteins complexes to connect the fusogenic transmembrane proteins with the cytoskeleton to enable myoblast fusion.

Although two distinct populations of myoblasts exist in Drosophila, this cellular diversity has not yet been validated in mammals. Here, we show that Tanc1 and Tanc2 are needed on both myoblast cells for proper fusion, like other essential vertebrate myoblast fusion components like Bai3, Dock1, Rac1, N-Wasp and the fusogenic protein Myomaker^29-31,40,41,52^. The potential crosstalk between Tanc1/2 and other known molecules that promote fusion should be investigated. While Ants/Rols7 interacts with Mbc in *Drosophila*^24^, we failed to detect this interaction with mammalian proteins (data not shown). It was proposed that Ants/Rols7 translocates to the fusion sites on exocytic vesicles^25^. While the mode of travel of Myomaker and Myomixer is unclear, they are likely to be transported on vesicles as membrane proteins. This could imply a potential cooperation between Tanc1/2 and Myomaker and/or Myomixer. The molecular mechanisms behind this hypothesis remain to be fully investigated.

We also demonstrated that the loss of fusion induced by either Tanc1 or Tanc2 depletion could be completely rescued by the overexpression of the reciprocal proteins. We concluded that Tanc1/2 carry out similar essential functions and that a threshold level of their expression is critical for efficient fusion. Interestingly, although Tanc1 knockout mice are viable^44^, the total knockout of Tanc2 in mice leads to lethality depending on the genetic background (https://www.mousephenotype.org/data/genes/MGI:2444121#phenotypesTab?dataSearch=weight)^53^. It would be interesting to evaluate the expression levels of Tanc2 in Tanc1 knockout mice. Also, while the phenotypes of these knockout mice have been well studied in the brain, their effect on skeletal muscles remains unknown. Generating new Tanc1 and Tanc2 conditional knockout mouse models would be necessary to establish the role of Tanc1/2 in myoblast fusion during embryogenesis and muscle repair.

Finally, we attempted to rescue the fusion defects in r*ols* mutant embryos by overexpressing human Tanc1 and Tanc2, using conditions in which Ants/Rols was previously shown to work. While we cannot rule out complications linked to differences in transgene expression levels, these assays suggested that individually, Tanc1 and Tanc2 could not replace the function of Ants/Rols7 in fusion. Potentially, expression of both Tanc1 and Tanc2 could be needed to rescue *rols7* fusion defects. Generating Tanc1 and Tanc2 co-expressing embryos would address this. There is some evidence for Kirrel1 and Kirrel3, two of the orthologs of the Kirre/Duf receptor that recruits Ants/Rols7 to the site of fusion in flies, participating in myoblast fusion in mammals^54,55^. While Tanc1 and Tanc2 may be recruited by these proteins during myoblast fusion in mammals, sequence differences may prevent them from interacting with Kirre/Duf in flies. Alternatively, there may be more fundamental mechanistic differences in fusion between flies and mammals that explain the lack of rescue. Nonetheless, this system could be of great value for future rescue experiments in the field of fusion.

## Material and Methods

### Antibodies

Mouse Monoclonal anti-Myosin Heavy Chain (MF20) (Immunofluorescence: 1:20, Western Blot: 1:10), mouse Monoclonal anti-Myogenin (F5D) (Western Blot: 1:10) and mouse monoclonal anti-Even-skipped (Eve) (2B8) (Immunofluorescence: 1:400) were obtained from Developmental Studies Hybridoma Bank. Mouse Monoclonal MyoD (Western Blot: 1:200) was obtained from BD Pharmingen (Cat. 554130). Rabbit Polyclonal anti-Tanc1 (Western Blot: 1: 2000) was obtained from Bethyl Laboratories (Cat. A303-019A). Mouse monoclonal anti-Myc 9E10 (cat. sc-40; immunofluorescence 1: 100; Western blot: 1:1000) and anti-HA tag (F-7) (Cat. sc-7392; Immunofluorescence: 1:100) and rabbit monoclonal anti-Tanc2 (Cat. sc-515710; Western Blot: 1: 20) were obtained from Santa Cruz Biotechnology Inc. Mouse monoclonal anti-Tubulin (1: 10,000) was obtained from Sigma-Aldrich (Cat. T5168). Rat Monoclonal anti-Myosin (MAC 147) (cross-reacts with Myosin in Drosophila; Immunofluorescence: 1:100) was obtained from Abcam (Cat. Ab51098). Rabbit anti-GFP (TP401) (Immunofluorescence: 1:100) was obtained from Torrey Pines Biolabs (Secaucus, NJ, USA).

### Cell culture, Transfection and C2C12 differentiation assay

C2C12 (ATCC) and Sol8 (Kind gift from Dr. Jacques Drouin, IRCM) murine myoblast cells were maintained in Dulbecco’s Modified Eagle Medium (DMEM) containing 20% FBS (vol/vol) and a mixture of antibiotics (penicillin and streptomycin (Gibco)). Transfection with 5 ug total DNA in a 6 well plate was performed using Lipofectamine 3000 (Invitrogen) according to the manufacturer’s protocol. ecoPhoenix retroviral packaging cells (kind gift of Dr. André Veillette (IRCM)) were maintained in DMEM containing 10% FBS supplemented with penicillin and streptomycin and were transfected in 10 cm plates with 10 ug of the indicated plasmid DNA at 50% confluency using the calcium phosphate precipitation protocol. Myc-tagged human Tanc1 and Tanc2 vectors were used for rescue experiments (kind gift from Dr. Stéphane Angers). Exponentially growing C2C12 and Sol8 (80-90% confluent) cells were differentiated by switching them into a media containing 2% horse serum and penicillin and streptomycin for 48 or 96 hours. A fusion index (%) was calculated from the ratio between the number of nuclei in Myhc-positive fibers containing 3 or more nuclei to the total number of nuclei in the same field multiplied by 100.

### ShRNA vectors to deplete Tanc1 and Tanc2

The pSIREN-RetroQ-ZsGreen retroviral system was used to generate the shRNA plasmids as previously described ^41^. Briefly, different shRNA oligonucleotides specifically targeting Tanc1 and Tanc2 were designed (please refer to Supplementary Table S1 for the list of oligo sequences). The pSIREN-RetroQ-ZsGFP vector was digested using BamHI/EcoRI and annealed primers were cloned into it. The constructs were verified by sequencing. The purified plasmids were transfected into the ecoPhoenix cells as described above. After 48h, the supernatants containing the indicated retroviruses were filtered and mixed with polybrene (Sigma) to a final concentration of 5 ug/ml. Two infections 24 h apart were performed on the C2C12 and Sol8 using the viral supernatants adjusted to 20% FBS. After 48 h, ZsGreen (GFP) positive myoblasts were sorted using a MOFLO cell sorter (Beckman Coulter). The efficiency of Tanc1 and Tanc2 knockdown was confirmed by real-time Q-RT-PCR as described next.

### Real-time polymerase chain reaction (Q-RT-PCR)

TRIZOL reagent (Invitrogen) was used to extract total RNA from the C2C12 and Sol8 cell lines infected with the indicated shRNAs, as recommended by the manufacturer. Dnase1 (Invitrogen) treatment on the total RNA was performed and cDNAs were generated using the M-MuLV Reverse Transcriptase and random primers (NEB), as per the manufacturer’s protocol. Real-time Q-RT-PCR system (ViiA7-96, Life technologies) was used to confirm specific knockdown of Tanc1 and Tanc2 by using SYBR Green PCR Master Mix (Applied Biosystems). Melt curve analysis was conducted for each primer set (See Supplementary Table S2 for the list of primers used) to confirm the specificity of the reaction. The beta2-microglobulin (B2m) gene was used as a control. The following protocol was used for all the real-time Q-RT-PCR reactions: 95 °C for 20 sec, followed by 40 cycles of 95 °C for 15 sec and 60 °C of 20 sec.

### Immunofluorescence and Microscopy of C2C12 cells

For overexpressed Myc-Tanc1 and Myc-Tanc2 localization, C2C12 cells were plated on fibronectin-coated glass coverslips. For coating, coverslips were incubated at 37°C with fibronectin (VWR, Cat. CACB356008) at a concentration of 5 ug/cm^2^ for 2 h. The coverslips were then rinsed with PBS twice and the cells were plated and incubated for 24 h prior to fixation, as described above. Cells were fixed for 15 min with 4% paraformaldehyde, washed with PBS and then permeabilized using 0.2% Triton X-100/ PBS solution for 5 min. Then, cells were incubated with anti-Myc (9E10) primary antibodies diluted 1:100 in wash buffer (TBS (200 mM Tris-HCL pH 7.6; 1.37 M NaCl), 1% BSA, 0.1% Triton) overnight at 4°C. Next, cells were washed 5 times with the wash buffer and then incubated at room temperature for 1h with Alexa 488-conjugated chicken anti-mouse secondary antibody together with Alexa-568-conjugated phalloidin (1:500) and Hoechst (1:10,000; Invitrogen). Cells were then washed 8 times with wash buffer and mounted using mowiol (Sigma, Cat#81381). The localization of Myc-Tanc1/2 was imaged using Zen Black software on a Zeiss LSM700 confocal microscope at a 63X magnification. The percentage of Myc+ cells presenting enrichment at the cell cortex or along stress fibers was estimated on 128 cells per conditions coming from 3 independent experiments.

For the effect of Tanc1 and Tanc2 knockdown on myoblast fusion, GFP-sorted C2C12 or Sol8 cells expressing scrambled versus Tanc1 or Tanc2 shRNA were plated on plastic and differentiated as described above. Cell fixation and immunostaining were performed as above except that permeabilization with 0.2% Triton was performed for 15min. Incubation with mouse monoclonal anti-MyHC (MF20; 1:20) was followed by incubation with Alexa 568-conjugated goat anti-mouse secondary antibody together with Hoechst. The effect of the Tanc1 and Tanc2 knockdown on the fusion phenotype of myoblasts was detected using the Leica DMIRB microscopy was used at an objective 20X magnification. All pictures were mounted using Fiji.

### Western blots

For protein expression, C2C12 myoblast cells were lysed using RIPA buffer (50 mM Tris pH 7.6; 0.1% SDS, 0.5% sodium deoxycholate, 1% NP-40, 5 mM EDTA, 150mM NaCl) for 30 min on ice. Protein concentration in the collected lysates was determined using the DC Reagent Assay kit from Bio-Rad as per the manufacturer’s protocol (Cat. 500-0116). 50ug of cleared extracts were denatured, fractionated on an 8% SDS-PAGE acrylamide gel, transferred on a nitrocellulose membrane (70 V for 2 h) and incubated in a 1% BSA/TBST blocking solution at RT for 1 h. After blocking, the membranes were incubated with the appropriate primary antibodies diluted in the blocking solutions at 4°C overnight. Unbound primary antibodies are then washed off with 1X TBS/0.1% Tween-20 solution and the membranes were incubated with the appropriate secondary antibodies for 1 h at RT. The specific signals were revealed using the Clarity Western ECL Substrate from Bio-Rad (Cat. 170-5061).

### Mixed population Assay

Parental Cells or cells expressing the indicated shRNA were labeled with red (PKH 26, Sigma) or green (PKH67, Sigma) lipid dyes according to manufacturer’s protocol, as previously described ^40^. Cells were mixed in a 1:1 ratio in 6-well plates. 24 h later cells were subjected to 48 h differentiation assays. Images were taken using the DMIRB microscopy at an objective 20X and analyzed using Fiji. The percentage of green, red or yellow fibers containing 1, 2 or 3 and more nuclei was quantified on 5 images per condition per experiments for three independent experiments.

### Drosophila embryo analyses

*w*^*1118*^ and *24B-Gal4* strains were obtained from the Bloomington Drosophila Stock Center (Bloomington, IN, USA). *rols*^*T192*^ and *rols*^*T627*^ strains were generously provided by Dr. E. Chen (UT Southwestern, TX, USA). To generate *UAS-HA-Tanc1* and *UAS-HA-Tanc2* transgenic strains, Tanc1 and Tanc2 coding sequences were inserted by Gateway cloning in frame with an N-terminal 3xHA tag in pTHW, a modified version of pUAST (Drosophila Genomics Resource Center, Indiana University, Bloomington, IN, USA)^56^. The resulting transgenes were injected into *w*^*1118*^ embryos and transgenic strains isolated by standard methods (BestGene Inc. Chino Hills, CA, USA). Both strains used in this study had second chromosome insertions. To examine embryonic Tanc expression in pre-fused or fused myoblasts (at stages 13 and 15, respectively), homozygous *24B-Gal4* virgins were crossed to *w*^*1118*^ (control) or homozygous *UAS-HA-Tanc1* or *UAS-HA-Tanc2* males. For rescue experiments, *24B-Gal4,rols*^*T627*^*/TM3,arm-GFP* virgins were crossed to *UAS-HA-Tanc1;rols*^*T192*^*/TM3,arm-GFP* or *UAS-HA-Tanc2;rols*^*T192*^*/TM3,arm-GFP* males. Flies were put into cages with yeasted apple-juice agar plates prior to egg-laying in order to acclimatize. To obtain stage 13 embryos, the egg-laying period lasted for 3 hours at 25°C and then the agar plates with eggs were transferred to 18°C for 14 hours. To obtain stage 15 embryos, the egg-laying period lasted for 4 hours at 25°C and then the agar plates with eggs were transferred to 18°C for 16 hours. Embryos were collected and rinsed and then washed twice with wash solution (0.7% w/v NaCl, 0.3% Tween-20). The chorions were removed by incubation in a 4% bleach solution for no more than 2 min. Embryos were subsequently fixed in 8 mL of a 1:1 solution of 4% paraformaldehyde in PBS:*n*-Heptane for 25 minutes. The paraformaldehyde layer was removed and replaced with 4 mL of methanol, and embryos were devitellinized with vigorous shaking. Embryos were re-hydrated in phosphate-buffered saline/ 0.3% Triton-X 100 (PBT) and blocked in PBT/0.5% bovine serum albumin (BBT) for 1 hour prior to addition and incubation with primary antibodies against Myhc or Eve, diluted in BBT, at 4°C overnight. For crosses involving the *TM3, arm-GFP* balancer, anti-GFP staining was performed to allow identification of embryos carrying the balancer. After four washes in PBT, embryos were incubated for two hours at room temperature with fluorescently-labelled secondary antibodies diluted 1:100 in BBT (Alexa 488-conjugated goat anti-rabbit IgG, Alexa 555-conjugated goat anti-rat IgG, Alexa 647-conjugated goat anti-mouse IgG; Invitrogen). After five final washes in PBT, embryos were incubated overnight in mounting medium (10% PBS, 90% glycerol, 0.2% *n*-propyl gallate) at 4°C to equilibrate. Embryos were mounted on glass slides and imaged using Zen Black software on a Zeiss LSM700 confocal microscope. Fluorescence signals were analyzed using ImageJ. Quantification of Eve-positive nuclei is based on the average number counted in three to four segments each from a total of twelve embryos, compiled from two independent experiments.

### Sequence alignments

The protein sequences of Drosophila Ants/Rols7 and human Tanc1 and Tanc2 were aligned using Clustal Omega (REF: DOI: <u>10.1093/nar/gkz268</u>). Sequences used were rolling pebbles, isoform B [Drosophila melanogaster], Accession: NP_729778.1; TANC1 isoform 1 [Homo sapiens], Accession: NP_203752; TANC2 isoform 1 [Homo sapiens], Accession: NP_079461.

## Acknowledgments

We would like to thank Dr. Viviane Tran for helpful discussions. We would like to thank the IRCM microscopy (Dr. D. Filion) and Flow cytometry (Dr. E. Massicotte) facilities for their expert assistance on this project. We thank Drs. Stéphane Angers (U. Toronto), Elizabeth Chen (UT Southwestern) and André Veillette (IRCM) for generous gifts of reagents. A.L. was supported by a Molson-Bombardier studentship from the IRCM Foundation. J-.F.C (grant #153065) and D.R.H. (grant #162109) were supported by grants from the Canadian Institute of Health Research (CIHR). J-.F.C. holds the TRANSAT chair in Breast Cancer Research.

## Data availability

All source data is available from the authors, upon reasonable request.

## Competing interests

The authors declare no competing interests.

## Supplementary informations

**Supplementary Figure S1.**
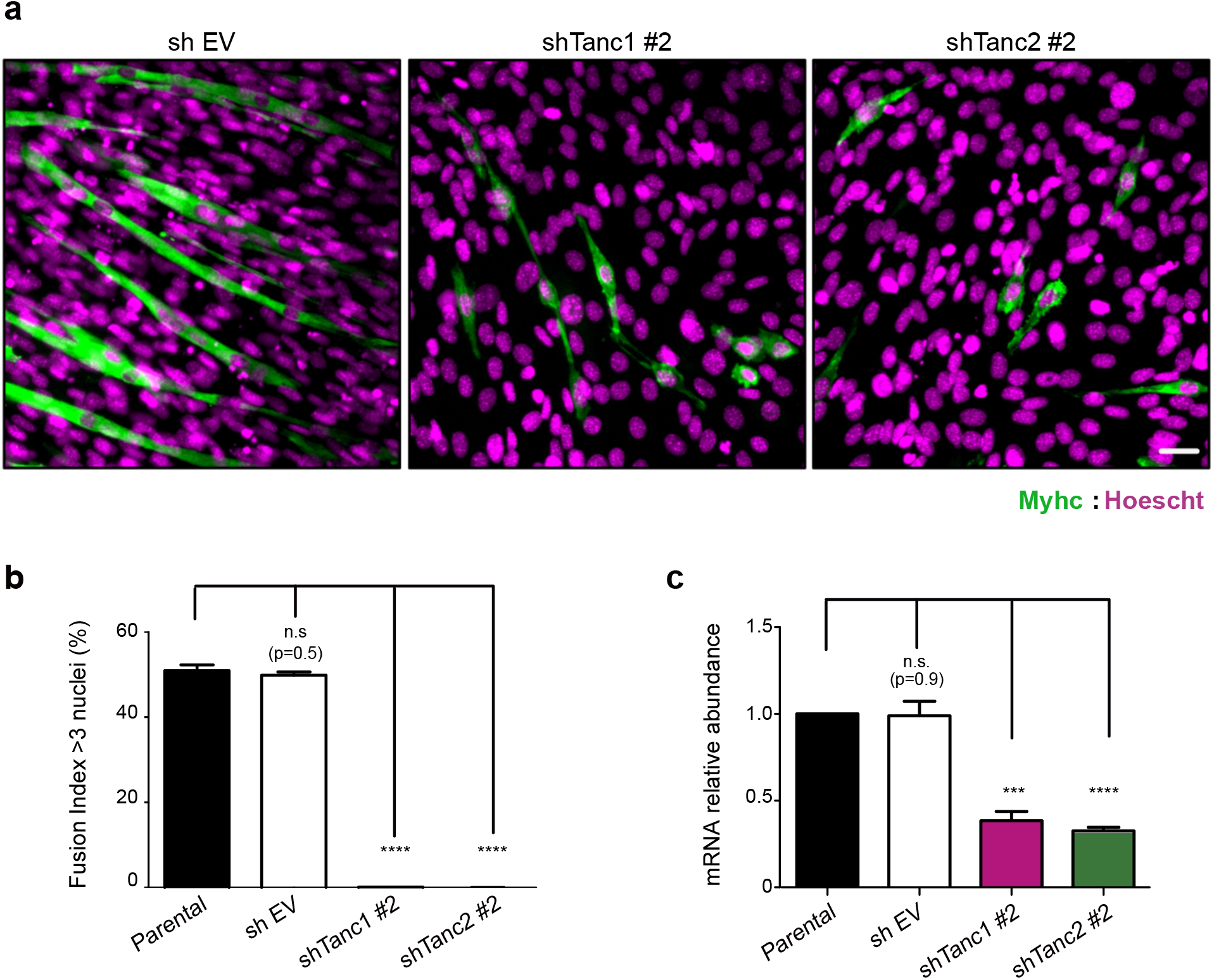
Depletion of Tanc1 or Tanc2 using a different set of shRNAs inhibits fusion. (a) C2C12 cells infected with shRNA sequence #2 against Tanc1 orTanc2 were enriched and submitted to differention for 72h. Fusion phenotypes of shTanc1#2 and shTanc2#2 was assed by microscopy after MHC (pseudocolored green) and nuclei (Hoescht, pseudocolored magenta) staining. No multinucleated fibers were detected in shTanc1#2 or shTanc2#2 conditions. (b) Fusion index representing the percentage of myofibers containing 3 or more nuclei was calculated on conditions presented in a. Data represent mean +/- SD of 3 independent experiments. (c) Validation of the specific knockdown of Tanc1 and Tanc2 by real-time Q-RT-PCR. The P value was calculated by one-way ANOVA followed by a Bonferroni test, ****P<0.0001. (Scale bar, 50 μm). Every experiment was replicated three times.

**Supplementary Figure S2.**
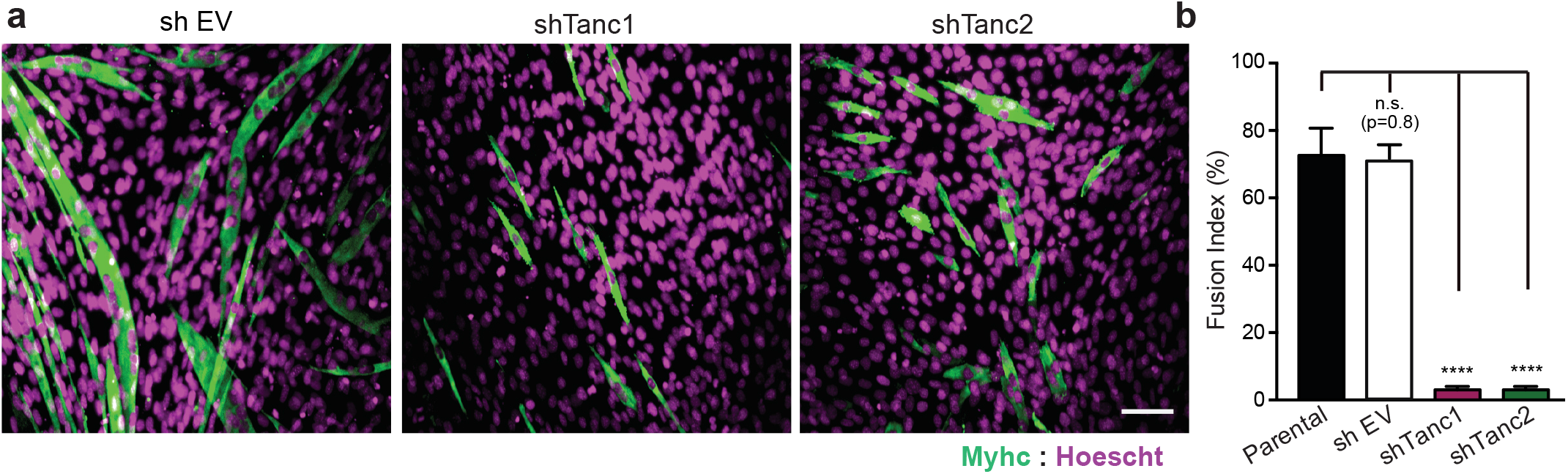
Depletion of Tanc1 or Tanc2 impairs myoblast fusion even after a long differentiation period. (a) Cells infected with an empty vector or vectors with shTanc1 or shTanc2 were stained with Myhc and Hoechst after 96 h to determine if fusion is blocked or delayed. No multinucleated fibers (>3 nuclei) were detected in shTanc1 and shTanc2. Scale bars, 50 μm (b) Fusion index was calculated on conditions presented in a. Data represent mean +/- SD of 3 independent experiments. The P value was calculated by one-way ANOVA followed by a Bonferroni test, ****P<0.0001. The experiment was replicated three times.

**Supplementary Figure S3.**
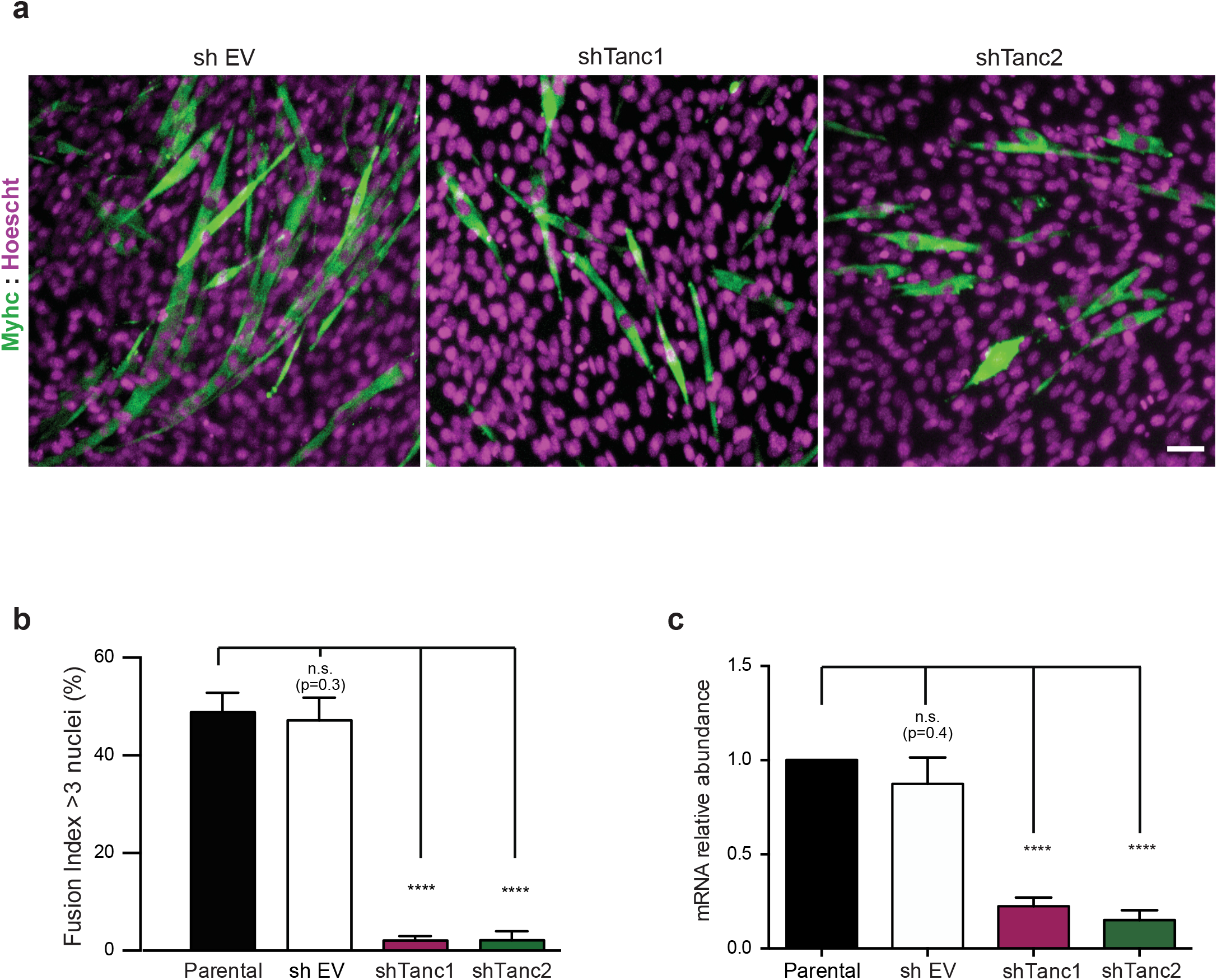
Tanc1 and Tanc2 are essential for myoblast fusion in a Sol8 myoblast cell system. (a-c). Sol8 myoblasts infected to express an empty vector or shRNAs against Tanc1 or Tanc2 were generated. These cells were differentiated and myofibers were stained for Myhc and Hoescht (nuclei). (a) Down-regulation of Tanc1 or Tanc2 blocks myoblast fusion after 96h of differentiation. (b) Fusion index was calculated on conditions presented in a. Data represent mean +/- SD of 3 independent experiments. (c) The specific knockdown of Tanc1 and Tanc2 in Sol8 was validated by real-time Q-RT-PCR. The P value was calculated by one-way ANOVA followed by a Bonferroni test, ****P<0.0001. Every experiment was replicated three times.

**Supplementary Figure S4.**
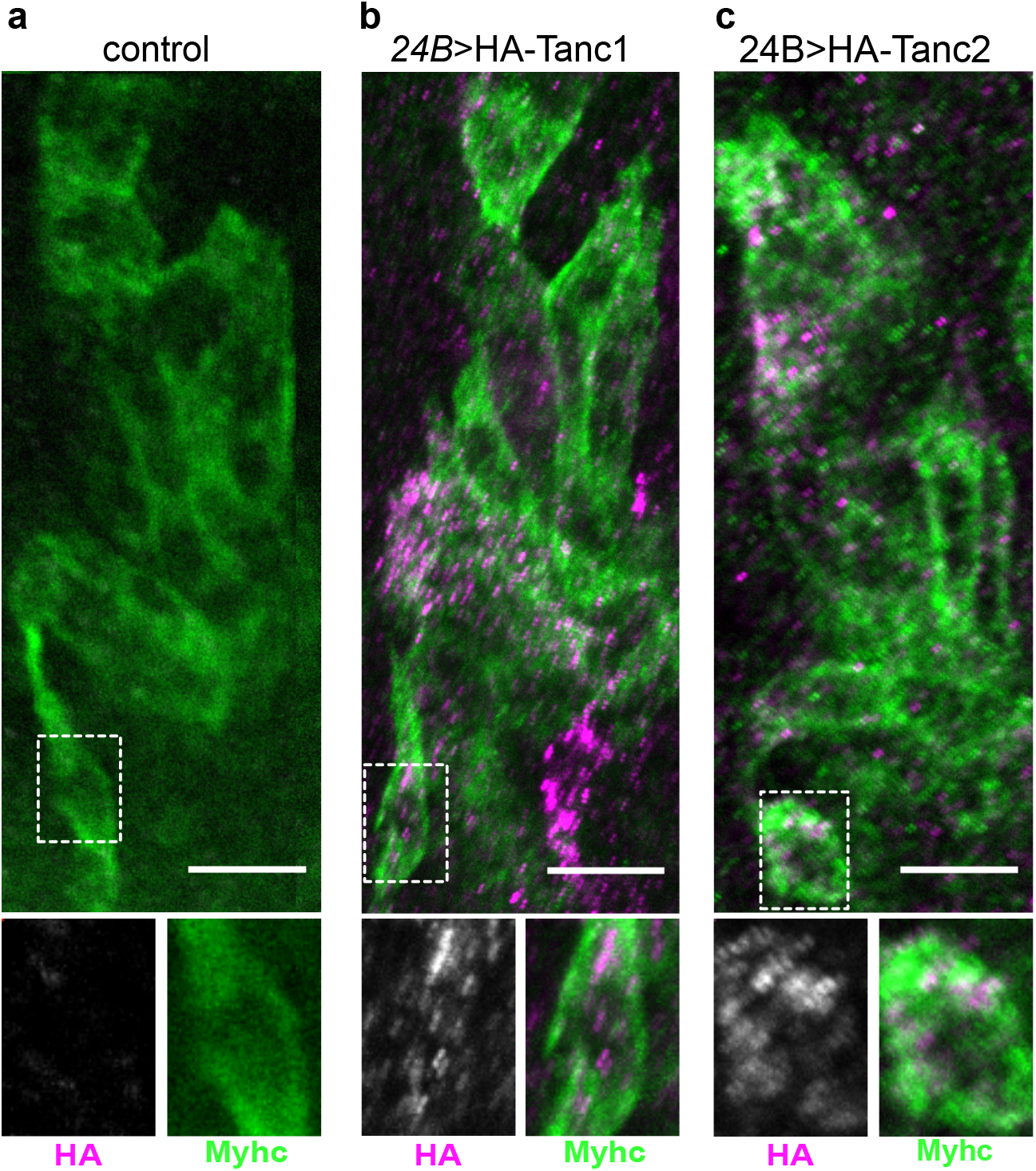
Transgenic HA-Tanc1 and HA-Tanc2 are expressed in pre-fusion myoblasts. (a-c) Detection of *24B-Gal4*-driven HA-Tanc1 and HA-Tanc2 expression in the developing musculature of stage 13 embryos, marked by Myhc staining. Both proteins were detected in individual myoblasts prior to fusion (insets). Scale bar: 10 μm. Genotypes: *w;24B-Gal4/+* (a); *w;UAS-HA-Tanc1/+;24B-Gal4/+* (b); *w;UAS-HA-Tanc2/+;24B-Gal4/+* (c).

**Supplementary Table S1.**
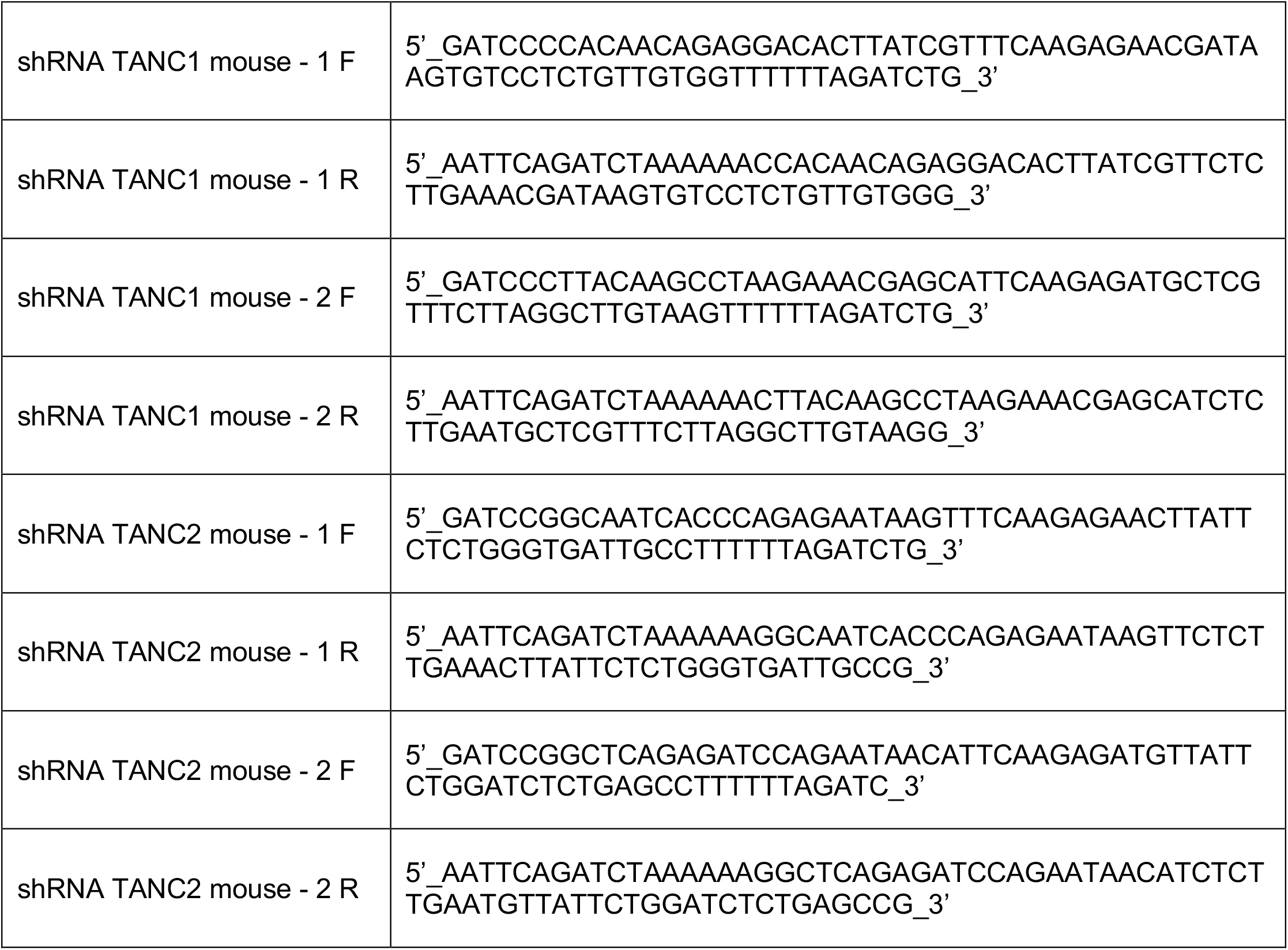
List of shRNA oligos

**Supplementary Table S2.**
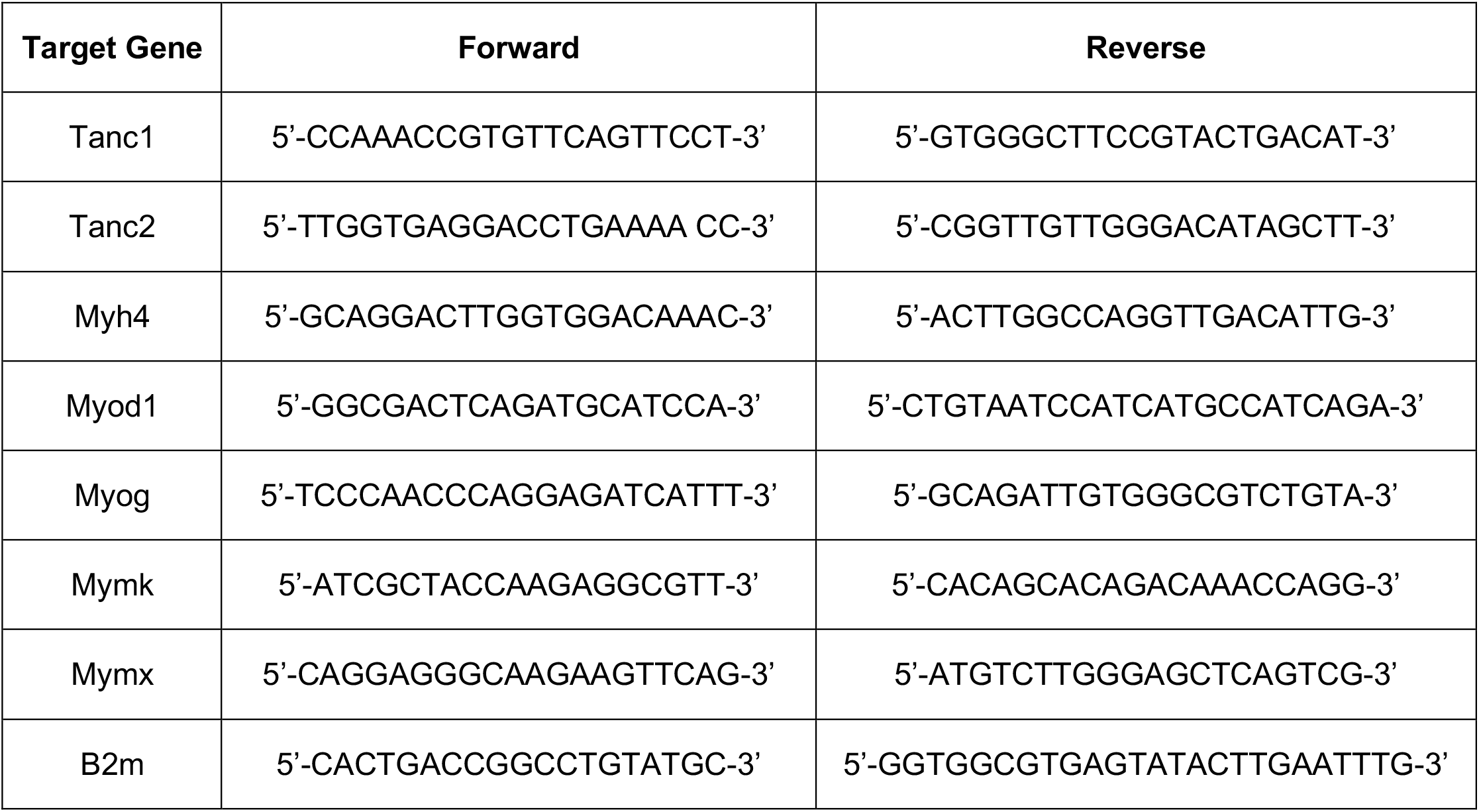
List of Q-RT-PCR primers.

## Notes

### Competing Interest Statement

The authors have declared no competing interest.

